# Requirement of microtubules for secretion of a micronemal protein CpTSP4 in the invasive stage of the apicomplexan *Cryptosporidium parvum*

**DOI:** 10.1101/2023.09.27.559857

**Authors:** Dongqiang Wang, Peng Jiang, Xiaodong Wu, Ying Zhang, Chenchen Wang, Meng Li, Mingxiao Liu, Jigang Yin, Guan Zhu

**Affiliations:** State Key Laboratory for Diagnosis and Treatment of Severe Zoonotic Infectious Diseases, Key Laboratory for Zoonosis Research of the Ministry of Education, Institute of Zoonosis, and College of Veterinary Medicine, Jilin University, Changchun 130062, China

**Author notes:** Correspondence (G.Z.).

**Keywords:** Protozoa, apicomplexan, *Cryptosporidium*, microneme, TSP-domain containing protein, secretion, microtubule, intracellular trafficking, heparin-binding domain/motif

## Abstract

The zoonotic *Cryptosporidium parvum* is a global contributor to infantile diarrheal diseases and opportunistic infections in immunocompromised/weakened individuals. Like other apicomplexans, it possesses several specialized secretory organelles, including micronemes, rhoptry and dense granules. However, the understanding of cryptosporidial micronemal composition and secretory pathway remains limited. Here we report a new micronemal protein in *C. parvum*, namely thrombospondin repeat domain-containing protein-4 (CpTSP4), providing insights into these ambiguities. Immunostaining and enzyme-linked assays show that CpTSP4 is prestored in the micronemes of unexcysted sporozoites but secreted during sporozoite excystation, gliding and invasion. In excysted sporozoites, CpTSP4 is also distributed on the two central microtubules unique to *Cryptosporidium*. The secretion and microtubular distribution could be completely blocked by selective kinesin-5 inhibitors, SB-743921 and SB-715992, resulting in the accumulation of CpTSP4 in micronemes. These support the kinesin-dependent microtubular trafficking of CpTSP4 for secretion. We also localize γ-tubulin, consistent with kinesin-dependent anterograde trafficking. Additionally, recombinant CpTSP4 displays nanomolar binding affinity to host cell surface, for which heparin acts as one of the host ligands. A novel heparin-binding motif is identified and biochemically validated for its contribution to the adhesive property of CpTSP4 by peptide competition assays and site-directed mutagenesis. These findings shed light on the mechanisms of intracellular trafficking and secretion of a cryptosporidial micronemal protein and the interaction of a TSP-family protein with host cells.

**Importance**

*Cryptosporidium parvum* is a globally distributed apicomplexan parasite infecting humans and/or animals. Like other apicomplexans, it possesses specialized secretory organelles in the zoites, in which micronemes discharge molecules to facilitate the movement and invasion of zoites. Although past and recent studies have identified several proteins in cryptosporidial micronemes, our understanding on the composition, secretory pathways and domain-ligand interactions of micronemal proteins remains limited. This study identifies a new micronemal protein, namely CpTSP4, that is discharged during excystation, gliding and invasion of *C. parvum* sporozoites. The CpTSP4 secretion depends on the intracellular trafficking on the two *Cryptosporidium*-unique microtubes that could be blocked by kinesin-5/Eg5 inhibitors. Additionally, a novel heparin-binding motif is identified and biochemically validated, which contributes to the nanomolar binding affinity of CpTSP4 to host cells. These findings indicate that kinesin-dependent microtubular trafficking is critical to CpTSP4 secretion and heparin/heparan sulfate is one of the ligands for this micronemal protein.

## Introduction

The zoonotic *Cryptosporidium parvum* is one of the leading causes of moderate to deadly diarrheal diseases in infants and opportunistic infections in immunocompromised patients (1–3). Like other apicomplexans (e.g., *Toxoplasma* and *Plasmodium*), cryptosporidial zoites possess several specialized secretory organelles, namely micronemes, rhoptries and dense granules (4, 5). Among them, micronemes are distributed in the sporozoite anterior and secrete various adhesins and peptidases to facilitate the moving zoites to cross the mucus layer and interact with host cells (5–8). Despite their importance in parasite invasion, only a limited number of micronemal proteins have been validated in *Cryptosporidium* (9–11). Studies on the secretory pathways of cryptosporidial micronemal proteins and their domain-ligand interactions are also limited. Mainly based on studies on *Toxoplasma* and *Plasmodium*, microneme secretion is triggered by a signaling cascade involving intracellular cyclic nucleotides, calcium level, phosphatidic acid and downstream effectors (6, 8). Micronemal proteins are thought to be secreted via membrane fusion of secretory vesicles with parasite plasma membrane, for which only a handful of molecular players have been identified (8, 12, 13). Cytoskeletons including microtubule and F-actin are known to participate in intracellular trafficking of vesicles in the secretory pathways (14, 15). *Toxoplasma* and *Plasmodium* possess an array of cortical/subpellicular microtubules (SPMTs) for maintaining the shape of zoites (16), while currently there is no evidence that SPMTs are involved in intracellular trafficking of secretory molecules. The two short intraconoidal microtubules, unseen in *Cryptosporidium*, are implied for transporting “microtubule-associated vesicles” to the apical tip during the rhoptry docking and exocytosis (6, 8, 11). The F-actin, which is part of the glideosome in gliding zoites and the moving junction formed during the parasite invasion, is implied in apical positioning of rhoptries (17). On the other hand, *C. parvum* lacks SPMTs, but possessing two central microtubules that are extended from the apex to the central and posterior regions of the zoites, respectively (18). The two central microtubules appear unique to *Cryptosporidium*, but their biological roles remain unclear.

Here we report the cellular and biochemical characterizations of *C. parvum* a thrombospondin-repeat domain-containing protein (aka thrombospondin-related adhesive/anonymous protein), namely CpTSP4. CpTSP4 is prestored in the micronemes of the sporozoites, but secreted during the excystation, gliding motility and invasion of sporozoites. During secretion, CpTSP4 is transported on the two central microtubules in sporozoites. The secretion and microtubular trafficking could be fully blocked by selective kinesin-5 inhibitors. Additionally, CpTSP4 possesses a novel heparin-binding motif (HBM) that contributes to its nanomolar binding affinity to host cells. These studies reveal a biological function to the cryptosporidial-unique microtubules in the intracellular trafficking of a micronemal protein and uncover one of the host cell ligand for an apicomplexan TSP-family adhesive protein.

## Results

### CpTSP4 is prestored in micronemes, but transported along the two central microtubules for secretion in sporozoites during and after excystation

The *C. parvum* genome encodes 12 TSP-family proteins (CpTSP1 to 12) (see Table S1 in the supplemental material) (19, 20). Among them, CpTSP1 (aka TRAP-C1) and CpTSP8 were localized to the apical region and/or surface in excysted sporozoites (20–22). This study focused on CpTSP4 (Gene ID: cgd8_150; Genbank: XP_625479) based on its moderate size (488 aa) and relatively simple domain architecture, i.e., an N-terminal signal peptide (SP; 25 aa), a single PAN/APPLE domain, and two TSP1-like repeats, but lacking a transmembrane domain (TMD) (Fig. 1A). To determine the subcellular location of CpTSP4 in the parasite (see Fig. 1B for illustration of sporozoite), an anti-CpTSP4 mAb was raised in-house against a short epitope (^89^KIKKADSWQEC^99^). A single band from sporozoite crude extract was recognized by this mAb in western blot analysis, in which the signals could be eliminated by preincubating antibody with the peptide immunogen (Fig. 1C), supporting the mAb′s specificity. In intact/unexcysted oocysts that were ruptured by freeze-and-thaw to allow access of antibody to sporozoites, anti-CpTSP4 mAb labeled the anterior region of sporozoites by immunofluorescence assay (IFA) that could be colocalized with a rabbit polyclonal antibody (pAb) against CpGP900 – a known microneme marker (Fig. 1D; Fig. S1) (23, 24). The staining pattern differs from that of the filamentous microtubules labeled by an affinity-purified rabbit pAb on *C. parvum* β-tubulin (CpTubB) (Fig. 1E) (18). These observations indicate that CpTSP4 is prestored in the micronemes in sporozoites before excystation.

**Fig. 1.**
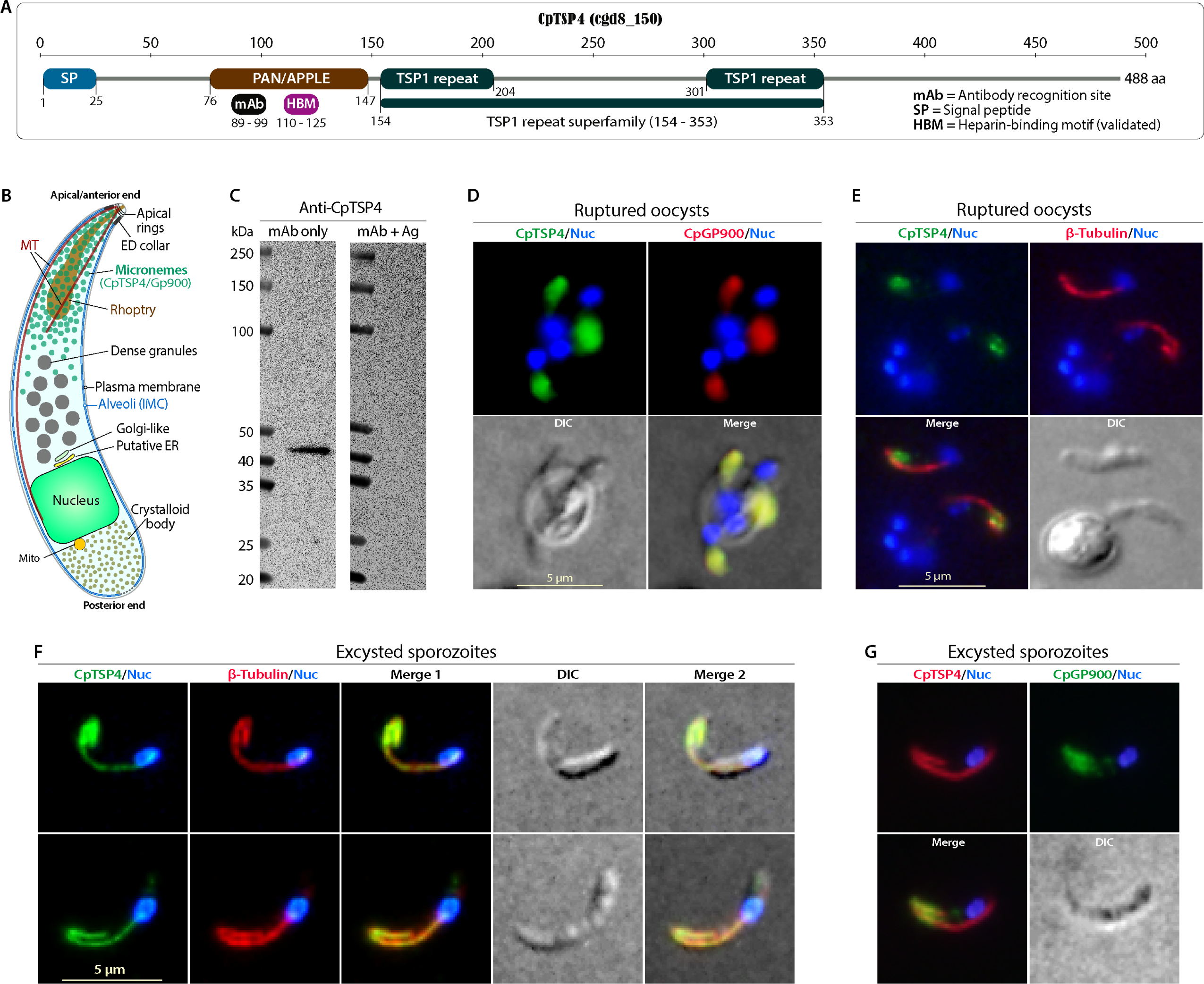
Distribution of CpTSP4 in the micronemes and on the microtubules in *C. parvum* sporozoites. (**A**) Architecture of CpTSP4 (cdg8_150) containing an N-terminal signal peptide (SP), a PAN/APPLE domain and two TSP1 repeats. mAb, the site recognized by the anti-CpTSP4 monoclonal antibody; HBM, heparin-binding motif determined in this study. (**B**) Illustration of the major structures in a *C. parvum* sporozoite, including micronemes and two unique central microtubules. ED collar, electron-dense collar; IMC, inner membrane complex; MT, microtubule. (**C**) Western blot detection of CpTSP4 in sporozoite crude extracts using anti-CpTSP4 mAb only (left), or the mAb pre-incubated with peptide antigen (mAb + Ag) (right). (**D – G**) Dual-labeling immunofluorescence assay (IFA) of CpTSP4 with CpGP900 (microneme marker) or microtubules in ruptured oocysts or excysted sporozoites, using an anti-CpTSP4 mAb, a rabbit anti-CpGP900 (C-terminal epitope) polyclonal antibody and a rabbit anti-CpTubB pAb. Intact oocysts were ruptured in fixative by three freeze-and-thaw cycles to allow access of antibodies to sporozoites. DIC, differential interference contrast microscopy; Nuc, nuclei counter-stained with 4,6-diamidino-2-phenylindole (DAPI).

Upon excystation, some CpTSP4 molecules started to show distribution on the two central microtubules that could be fully colocalized with anti-CpTubB pAb (Fig. 1F; Fig. S2), implying intracellular trafficking of CpTSP4 on the two microtubules. These microtubules were recently reported to be *Cryptosporidium*-unique (18), differing from SPMTs in *Toxoplasma* and *Plasmodium* (4, 25). The labeling of microtubules by anti-CpTSP4 mAb was not caused by its cross-reaction with *C. parvum* tubulin, because: 1) the mAb did not label the microtubules in unexcysted sporozoites (Fig. 1E); and 2) when anti-CpTSP4 mAb are anti-CpTubB pAb were individually pre-incubated with their peptide immunogens, each immunogen only eliminated fluorescent signals from corresponding antibody, but not those from the other antibody (see Fig. S3 in the supplemental material). For comparison, anti-CpGP900 pAb mainly labeled the micronemes in the anterior of sporozoites, showing no filamentous distribution (Fig. 1G). The labeling pattern of CpGP900 in excysted sporozoites is consistent with our previous study (23), implying that CpTSP4 and CpGP900 use different pathways for secretion.

**Fig. 2.**
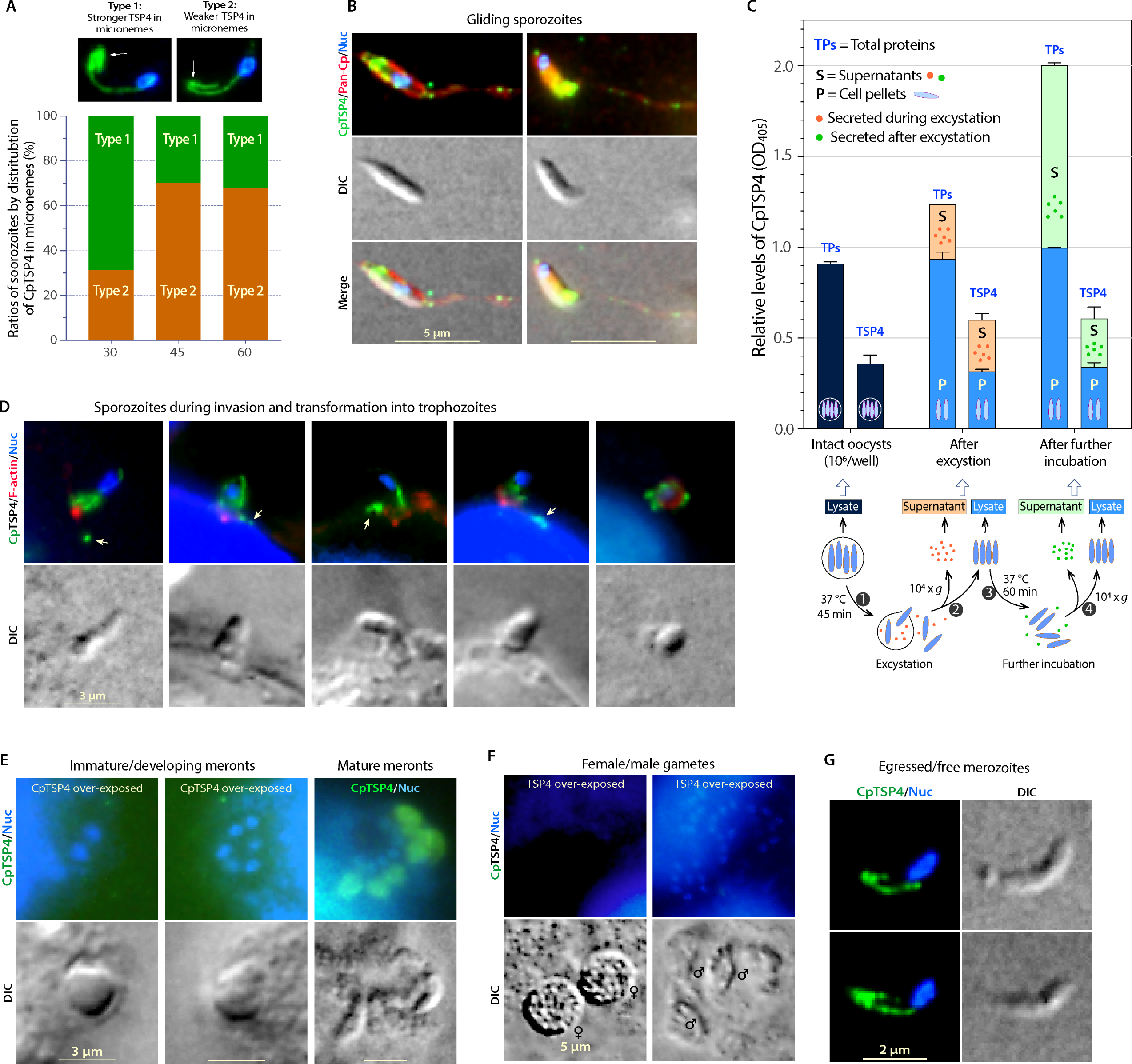
Distribution and secretion of CpTSP4 in various stages of *C. parvum* as detected with anti-CpTSP4 mAb. (**A**) During the course of excystation, the ratios of sporozoites showing strong CpTSP4 signals in the anterior region (marked as type I sporozoites) were reduced, whereas those showing weak anterior CpTSP4 signals (type II) were increased (N >200 in each time point). (**B**) Immunostaining of CpTSP4 (green) in free sporozoites allowed to glide on poly-L-lysine-coated slides in RPMI-1640 medium for 20 min at 37°C. Specimens were colocalized with rabbit antibody against total sporozoites proteins (marked as Pan-Cp) to better visualize the gliding trails (red). (**C**) Semi-quantitative ELISA detection of secreted CpTSP4 in the supernatants (S) and non-secreted CpTSP4 in the parasite cell pellets (P) collected immediately after excystation or after further incubation of excysted sporozoites in fresh RPMI-1640 medium. Intact oocysts were used as loading control. Lysates and supernatants were coated in microplates and detected with anti-CpTSP4 mAb. A rabbit pAb against total sporozoites proteins (TPs) were used in parallel for reference. Each samples contained 10^6^ oocysts or equivalent (N = 3/sample). Bars showed standard error of the mean (SEM). (**D**) Immunostaining of CpTSP4 (green) in sporozoites during the invasion and transformation into trophozoites. Host cell F-actin was stained with rhodamine-phalloidin (red). Arrows indicate CpTSP4 discharged into extracellular space. (**E**) Immunostaining of CpTSP4 in meronts, showing CpTSP4 is absent in immature meronts, but present in mature meronts containing developed merozoites. (**F**) Immunostaining of CpTSP4 in *C. parvum* sexual stages, showing absence of CpTSP4 in both male and female gametes. (**G**) Immunostaining of CpTSP4 in egressed merozoites, showing similar distribution of as seen in excysted sporozoites. DIC, differential interference contrast microscopy; Nuc, nuclei counter-stained with 4,6-diamidino-2-phenylindole (DAPI).

The secretion of CpTSP4 resulted in the reduction of CpTSP4 in the anterior of the sporozoites during the cause of excystation, while in the meantime the signals over the two microtubules remained relatively constant (Fig. 2A). Secreted CpTSP4 molecules were detected by IFA along the sporozoite gliding trails, mostly in granular form (Fig. 2B; Fig. S4). Semi-quantitative ELISA detected CpTSP4 from the supernatants collected after excystation (indicating secretion during excystation) and the supernatants when excysted sporozoites after washes were further incubated (indicating secretion after excystation) (Fig. 2C). The contents of intracellular CpTSP4 in unexcysted oocysts, excysted sporozoites, and further incubated sporozoites remained relatively constant. For comparison, a pAb against parasite total proteins detected similar patterns of secreted proteins (Fig. 2C). The data indicate that CpTSP4 is continuously secreted, and new CpTSP4 molecules are synthesized during and after excystation.

During the sporozoite invasion of host cells, CpTSP4 is continuously distributed over the shortening microtubules in the transforming sporozoites and discharged to the extracellular space (Fig. 2D). During intracellular development, CpTSP4 is absent in immature/developing meronts (2–6 nuclei), but present in mature/developed meronts containing fully developed merozoites (8 nuclei) (Fig. 2E; Fig. S5). The distribution on one side of nucleus in the packed meronts is similar to CpGP900 that is present only in fully developed merozoites (23). Also like CpGP900, CpTSP4 was absent in male and female gametes (Fig. 2F). In egressed merozoites, CpTSP4 was also localized to the anterior region and the two microtubules (Fig. 2G; Fig. S6), suggesting that CpTSP4 plays roles shared by sporozoites and merozoites. Collectively, CpTSP4 is a micronemal protein existing only in the parasite zoites. It is prestored in the oocysts, the environmental stage, and transported over the two central microtubular filaments for secretion from excysting and invading sporozoites. It is worth noting that, in agreement with the lack of TMD, CpTSP4 is absent on the sporozoite plasma membranes, suggesting that CpTSP4 is not attached to sporozoites via glycosylphosphatidylinositol (GPI) anchor or binding to other surface proteins in the parasite.

### Selective kinesin-5/Eg5 inhibitors (SB-743921 and SB-715992) could completely block the secretion of CpTSP4 and its transport on the central microtubules

Dynein and kinesin are the two molecular motors involved in microtubule trafficking (15, 26). To determine which motor was involved in microtubule trafficking of CpTSP4, two dynein inhibitors and two kinesin inhibitors (100 μM) were first tested for effects on the secretion of CpTSP4 during excystation. Strikingly, the secretion of CpTSP4 from sporozoites was completely suppressed by SB-743921 (CAS 940929-33-9), which is a highly selective inhibitor of human kinesin-5 (aka Eg5, or kinesin spindle protein [KSP]) (27, 28), but not by the other three inhibitors (Fig. 3A). The secretion of CpTSP4 from sporozoites could also be blocked by another kinesin-5/Eg5 inhibitor SB-715992 (CAS 336113-53-2; an SB-743921 analog) (Fig. 3B). SB-743921 inhibited the CpTSP4 secretion in a dose-dependent manner, showing low micromolar inhibitory activity (*EC*_50_ = 4.66 μM) (Fig. 3C). The inhibition of CpTSP4 secretion and transport on microtubules by SB-743921 was further confirmed by IFA (Fig. 3D; Fig. S7), in which the presence of SB-743921 (50 μM) during excystation resulted in the accumulation of CpTSP4 in the anterior region of the sporozoites and the loss of CpTSP4 signals from the two central microtubules. Under the same condition, SB-743921 had no apparent effect on the parasite morphology and excystation rate (Fig. 3D, 3E).

**Fig. 3.**
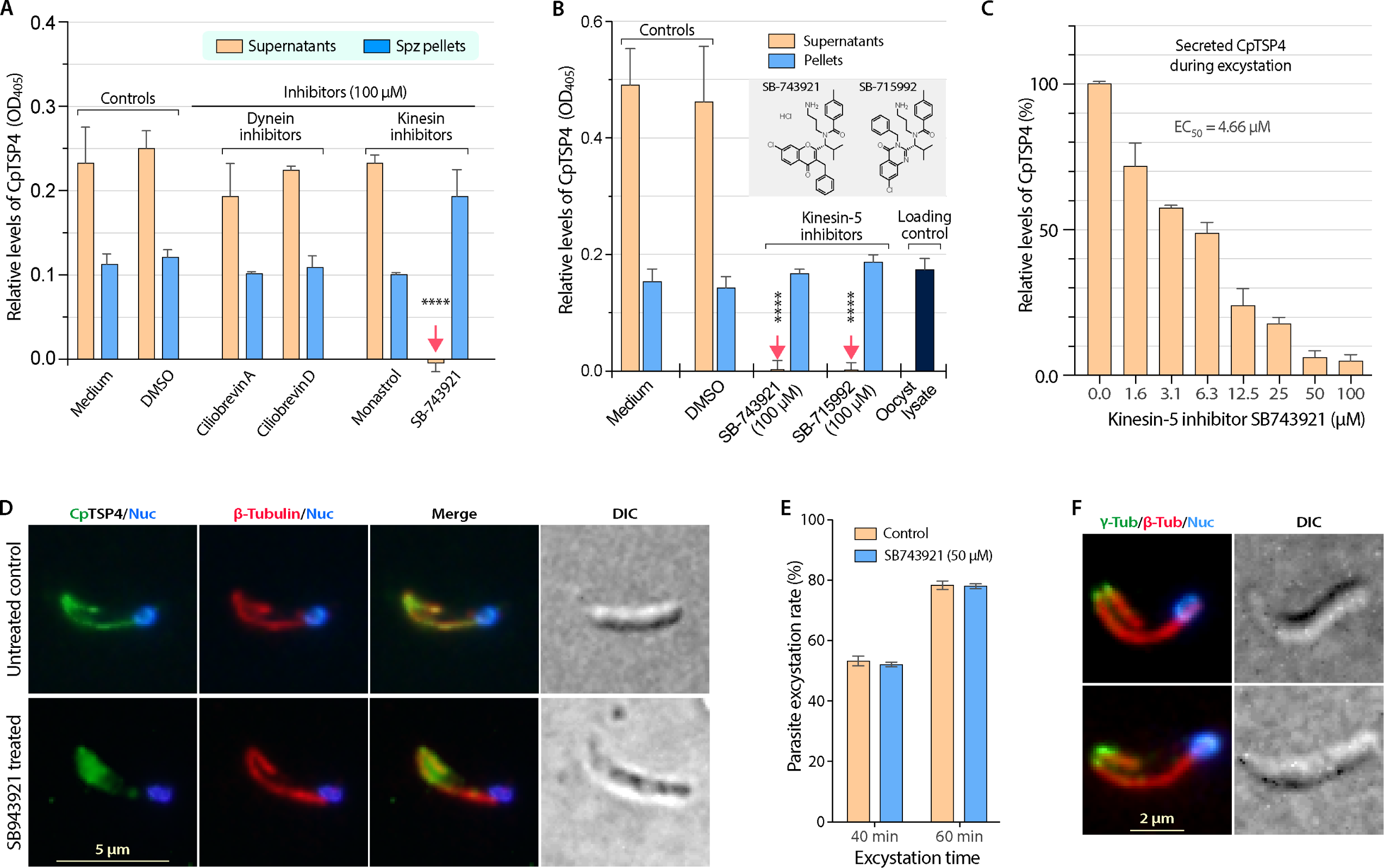
Effect of the selective kinesin-5 inhibitors SB-743921 and SB-715992 on the secretion of CpTSP4 and distribution in sporozoites. (**A**) Effects of selected dynein and kinesin inhibitors on the secretion of CpTSP4 from *C. parvum* during excystation. Oocysts (10^6^ per sample) were incubated in excystation medium with or without inhibitors (100 μM) for 1 h. Supernatants and lysates of sporozoite (Spz) pellets were coated in microplates, and CpTSP4 was detected by semi-quantitative ELISA using anti-CpTSP4 mAb. The selective kinesin-5 inhibitor SB-743921 completely inhibited the secretion of CpTSP4 (indicated by red arrows). (**B**) Complete inhibition of a second kinesin-5 inhibitor SB-715992 in comparison with SB-743921 (both at 100 μM; inset depicts their chemical structures) on the secretion of CpTSP4 from the parasite after excystation for 1 h by ELISA (10^6^ oocysts/sample). (**C**) Dose-dependent inhibition of SB-743921 on the secretion of CpTSP4 from the parasite after excystation for 1 h by ELISA (10^6^ oocysts/sample). (**D – E**) Treatment of SB-743921 at 50 μM resulted in the accumulation of CpTSP4 in the anterior region of excysting sporozoites and loss of its distribution on the two microtubules, but had no apparent effect on the parasite morphology (**D**) or on the parasite excystation rate (**E**). (**F**) Co-localization IFA of microtubules (red) using anti-CpTubB pAb with γ-tubulin (green) using a commercial anti-γ-tubulin mAb. In bar charts, error bars show standard error of the mean (SEM; N=3). **** = *P* <0.001 based on two-way ANOVA with Tukey’s multiple comparisons.

These observations suggests the essentialness of kinesin-dependent microtubule trafficking in the secretion of CpTSP4, likely a kinesin-5/Eg5 ortholog with reasonable homology at the motor domain (MD), the inhibitor-binding site. Indeed, the *C. parvum* genome encodes a unique kinesin-5 ortholog (designated as CpKin5; cgd6_4210). CpKin5-MD shares low homology to the other six *C. parvum* kinesins (similarity scores between 31.3% and 46.3%) (Table S2 in the supplemental material) (29), but higher homology to human kinesin-5/Eg5 (73.3% similarity; vs. 49.9% and 54.4% to the kinesin-5 orthologs from *Toxoplasma* and *Plasmodium*) (Table S3 and Fig. S8 in the supplemental material). Molecular docking also shows that CpKin5 and human kinesin-5/Eg5 share a conserved SB-743921-binding pocket (Fig. S9 in the supplemental material) (28). While more experimental evidences are needed, the current data strongly suggest that CpKin5 is the likely molecular motor involved in the microtubular transport of CpTSP4.

The involvement of kinesin suggests an anterograde transport of CpTSP4-cargoes towards the posterior end on the two central microtubules (i.e., from the minus to the plus end), in which the minus end is typically attached to microtubule organization center (MTOC). In *Toxoplasma* and *Plasmodium*, apical polar rings serve as unconventional MTOC for SPMTs. However, we were able to detect γ-tubulin at the anterior ends of the two central microtubules (Fig. 3F), suggesting that *C. parvum* may use conventional γ-tubulin-based MTOC for these microtubules. The polarity of the central microtubules agrees with the function of kinesins that slide towards the plus end of microtubules for transporting CpTSP4-cargoes from the anterior micronemes.

### A novel heparin-binding motif contributes to approximately half of the host cell-binding activity of CpTSP4

The binding property of CpTSP4 on host cells was investigated to gain insight into its biological roles using GST-fused recombinant CpTSP4 (rCpTSP4; excluding signal peptide) (Fig. 4A, 4B). rCpTSP4 could specifically bind to formalin-fixed HCT-8 cell monolayers as determined by flowcytometry and IFA (vs. no binding by GST-tag) (Fig. 4C, 4D). The binding followed a one-site specific binding kinetics, showing nanomolar binding affinity (App. *K*_d_ = 334 nM) (Fig. 4D).

**Fig. 4.**
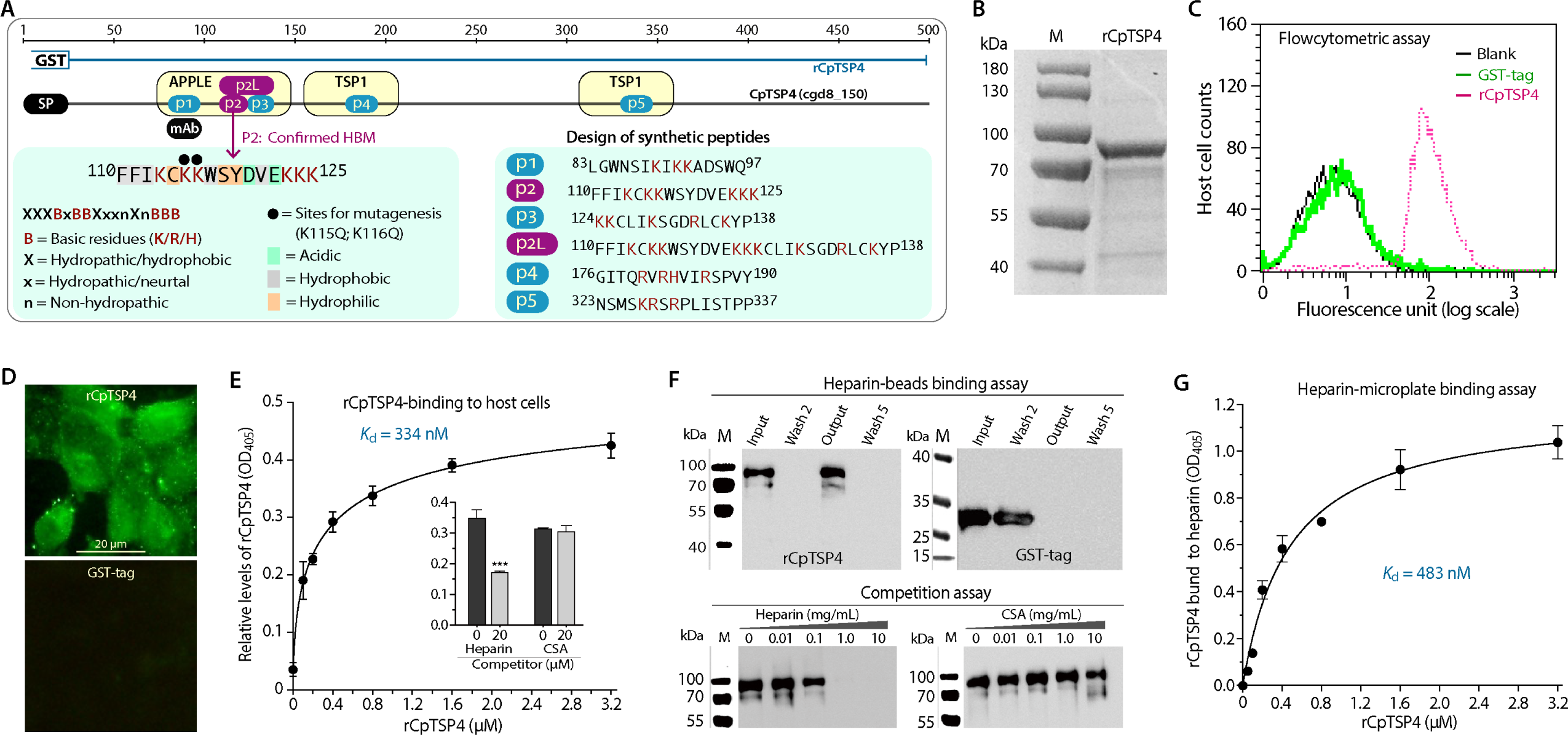
Specific binding of GST-fused recombinant CpTSP4 (rCpTSP4) to host cells and to heparin. (**A**) Illustration of the positions of CpTSP4 domains, rCpTSP4, candidate heparin-binding motif (HBM; sites and sequences) for designing synthetic peptides (p1 to p5), and the two amino acids at the p2 site for site-directed mutagenesis (K115Q/K116Q). Purple color indicates the novel HBM site determined in this study. (**B**) Purified rCpTSP4 fractionated by SDS-PAGE. (**C–D**) Binding of rCpTSP4 (vs. GST-tag) to host cells as determined by flowcytometry (**C**) and by immunofluorescence-based assay (**D**). (**E**) Binding kinetics of rCpTSP4 to formalin-fixed HCT-8 cells by enzyme-linked assay (App. *K*_d_ = 334 nM). Inset shows that the binding of rCpTSP4 (2.0 μM) to HCT-8 cells could be reduced by heparin (20 μM) but not by chondroitin sulfate A (CSA; 20 μM). *** = *P*<0.01 by two-tailed Student’s *t*-test. (**F**) Binding of rCpTSP4 to heparin-agarose beads (vs. GST-tag; upper panel) and competition of the binding by heparin (vs. CSA; lower panel). In binding assay, heparin-beads were mixed with rCpTSP4 or GST and washed five times. Original solution (Input), selected flowthroughs (Wash 2 and Wash 5) and the beads after washes (Output) were fractionated in SDS-PAGE and analyzed by western blotting. In competition assay, heparin-beads were incubated with rCpTSP4 in the presence of heparin or CSA at specified concentrations, washed and analyzed by western blotting. (**G**) Binding kinetics of rCpTSP4 to heparin by enzyme-linked assay (App. *K*_d_ = 483 nM), in which heparin was coated in microplates for binding by rCpTSP4 and detection by ELISA-like procedures. This assay used GST-tag at the same molar concentrations as negative control and for background signal subtraction. In **C** to **G**, GST-fused rCpTSP4 and GST-tag were detected using an anti-GST mAb and secondary antibodies conjugated with Alex Fluor-488 (**C** and **D**) or alkaline phosphatase (**E** to **G**). Errors bars in **E** and **G** show standard error of the mean (SEM; N=3).

CpTSP4 contains four regions rich in basic amino acids (BAAs), which is a common feature for heparin-binding motifs (HBMs) (Fig. 4A) (30, 31). We hence hypothesized that heparin/heparan sulfate (HS) might serve as the ligand (or one of the ligands) for CpTSP4. To test the hypothesis, we first performed a competition assay, showing that heparin, but not the closely related glycosaminoglycan chondroitin sulfate A (CSA), could compete with rCpTSP4 for the host cell-binding activity (Fig. 4E, inset). Second, direct binding of rCpTSP4 to heparin-conjugated agarose beads was observed in column-based assay (vs. no binding by GTS-tag) (Fig. 4F; upper panel). The binding of rCpTSP4 to heparin-beads could be specifically competed out by heparin, but not by CSA (Fig. 4F; lower panel). Third, using an enzyme-linked microplate assay developed in this study, in which the microplates were coated with heparin (10 μM) and the binding of rCpTSP4 (vs. GST-tag) were detected using anti-GST mAb, we observed a one-site specific binding kinetics of GST-CpTSP4 to heparin with sub-micromolar binding affinity (App. *K*_d_ = 483 nM) (Fig. 4G).

We then proceeded to identify the HBM in CpTSP4. Six peptides corresponding to the four BAA-rich sites in CpTSP4 were synthesized (see sequences and positions in Fig. 4A) and used in competition and binding assays. Site #2 was located in a PAN/APPLE domain and contained a relatively long BAA-rich sequence, for which three peptides were synthesized, including a long peptide (p2L) and two short ones (p2 and p3). In microplate-based competition assay, p2 and p2L peptides reduced the binding of rCpTSP4 to heparin in a dose-dependent manner (Fig. 5A), indicating that the p2 sequence was the HBM. Direct binding of p2-peptide to heparin was also confirmed, in which fluorescein isothiocyanate (FITC)-conjugated p2-peptide displayed low-micromolar binding kinetics to heparin in microplate-based fluorescence assay (App. *K*_d_ = 3.39 μM) (Fig. 5B). FITC-p2 also displayed low-micromolar binding kinetics to host cells (App. *K*_d_ = 6.23 and 7.03 μM) to HCT-8 and MDBK cells, respectively (Fig. 5C). Additionally, p2-peptide (2 μM) could compete with rCpTSP4 (1 μM) for binding to HCT-8 cells (vs. no competition by p1-peptide used as a control) (Fig. 5D).

**Fig. 5.**
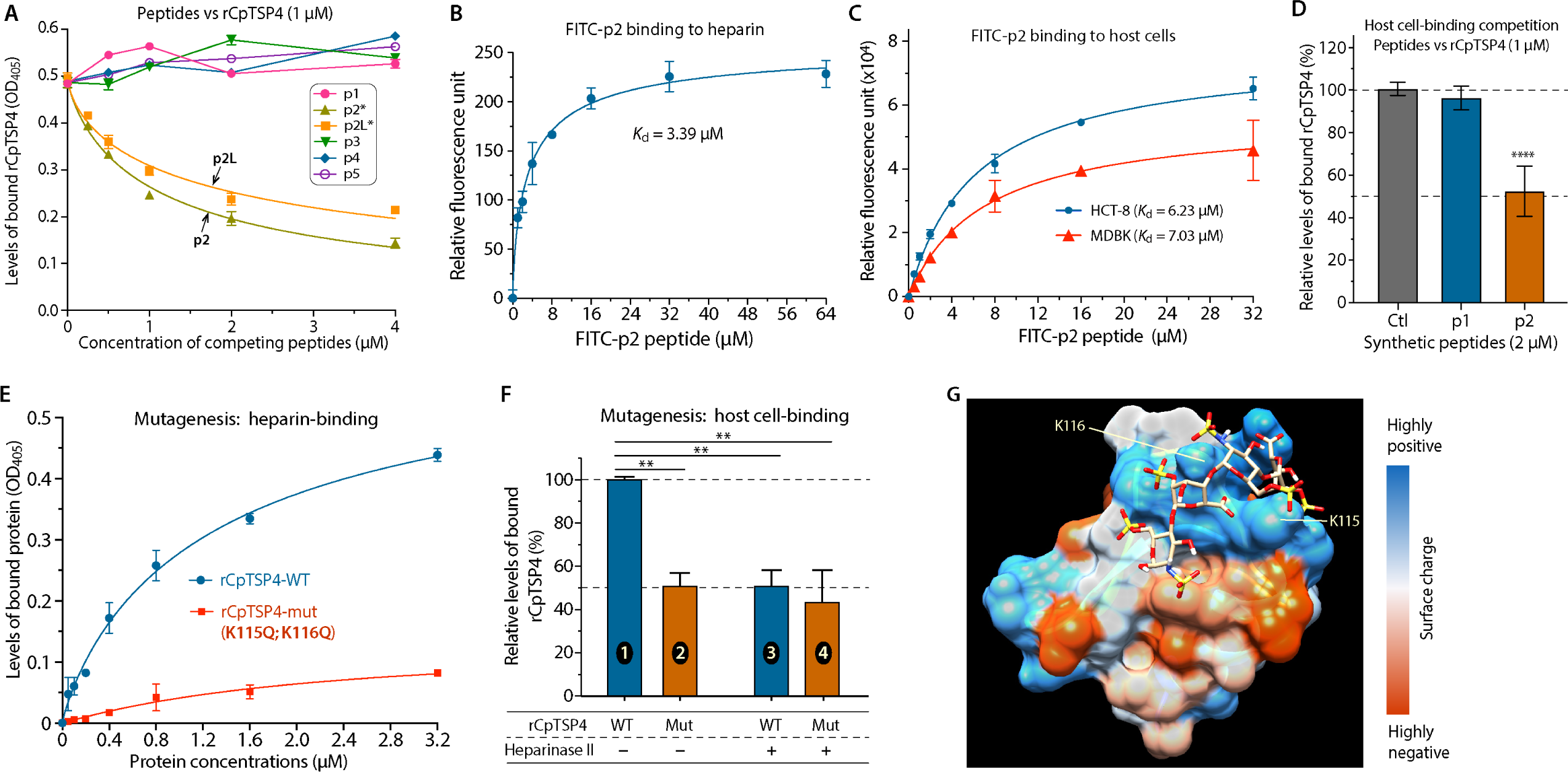
Determination of a novel heparin-binding motif (HBM) in CpTSP4 by peptide competition assay and site-directed mutagenesis. (**A**) Competition assay showing that the binding of rCpTSP4 to heparin coated in microplates could be competed out by p2 and p2L peptides in a dose-dependent manner, but not by other four synthetic peptides. Bound rCpTSP4 was detected by anti-GST mAb similar to ELISA. (**B**) Binding kinetics of FITC-conjugated p2 peptide to heparin coated in microplates (App. *K*_d_ = 3.39 μM). (**C**) Binding kinetics of FITC-conjugated p2-peptide to formalin-fixed HCT-8 and MDBK cells (App. *K*_d_ = 6.23 and 7.03 μM, respectively). (**D**) The binding rCpTSP4 (1.0 μM) to fixed HCT-8 cells was reduced by p2-peptide (2.0 μM) by 47.9% (vs. 4.2% by the control p1-peptide). **** = *P*<0.001 by Sidak multiple comparison test. (**E**) Comparison of the heparin-binding kinetics between wild-type and mutated rCpTSP4 (rCpTSP4-WT vs. rCpTSP4-mut), showing that the mutation of two Lys residues at the p2 site (K115Q/K116Q) eliminated the specific binding of rCpTSP4 to heparin in the enzyme-linked microplate assay. (**F**) The mutations (K115Q/K116Q) reduced the binding of WT rCpTSP4 to HCT-8 cells by 49.3% (bars 1 vs. 2), similar to the 49.1% reduction caused by heparinase II treatment of host cells (bars 1 vs. 3). Mutations at the p2-site (bars 2 and 4) or heparinase II treatment of host cells (bars 3 and 4) produced similar effect on the binding of rCpTSP4. ** = *P*<0.1 by Sidak multiple comparison test. (**G**) Molecular docking showing the binding of a tetrameric heparin to a positively charged crevice at the p2 site of CpTSP4, in which the Lys residues subjected to mutagenesis were indicated. Errors bars = standard error of the mean (SEM; N=3).

To further validate that the p2 sequence (^110^FFIKCKKWSYDVEKKK^125^) was responsible for the heparin-binding of CpTSP4, two Lys residues with a polar basic side chain at positions 115 and 116 were replaced by two Gln residues with a polar neutral side chain (i.e., K115Q and K116Q) by site-directed mutagenesis (see illustration in Fig. 4A). In microplate assay, the mutation essentially abolished the binding of rCpTSP4 to heparin (Fig. 5E). There was a low level of non-specific binding of mutated rCpTSP4, though (Fig. 5E, red line), which was likely attributed to the electrostatic interactions between the clustered basic residues in the protein and negatively charged heparin. In host cell-binding assay, the mutations reduced the binding of rCpTSP4 to HCT-8 cells by ∼50% (Fig. 5F, bars 1 vs 2). Pre-digestion of HCT-8 cells with heparinase II also reduced the binding of rCpTSP4 to HCT-8 cells by ∼50%, but had no significant effect on the binding of mutated protein (that was already ∼50% lower than the wild-type rCpTSP4) (Fig. 5F, bars 3 vs 4). We noticed that, under the physiological salt concentration (150 mM NaCl), non-specific binding of mutated rCpTSP4 to heparin or host cells was relatively high. The issue was resolved using high-salt buffer (PBS containing 500 mM NaCl and 1% Tween-20; see more details in Materials and Methods). High-salt buffer was used in all binding assays in this study to minimize the non-specific binding of rCpTSP4 to heparin or host cells. Heparin-binding at the p2 site was also supported by molecular docking with a structural model comprised of 51 amino acids spanning the p2 site (amino acid positions 50 to 150) built by AlphaFold (32). The prediction of heparin-binding used ClusPro server (https://cluspro.bu.edu/) under the preset “heparin-binding mode” (33), in which the heparin tetramer could fit well into a positively charged crevice at the p2 site (Fig. 5G).

The discovery that CpTSP4 possesses heparin-binding property expands the concept of heparin-binding proteins to include a protozoan TSP protein. The amino acid profile at the p2 site (XXXBxBBXxxnXnBBB; see Fig. 4A for annotation) also represents a novel HBM because it differs from known HBM consensus sequences (e.g., XBBXBX, XBBBXXBX, XBBBXXBBBXXBBX or TXXBXXTBXXXTBB; B=basic, X=hydropathic, T=turn) (30, 31). Heparin/HS are present in the intestinal mucus and extracellular matrix (ECM), and commonly explored as a host ligand by microorganisms including protozoan parasites for attachment (34–36). Based on the peptide competition assay and binding kinetics of wild-type and mutated CpTSP4 (Fig. 5), the p2 site HBM might contributes roughly 50% of the binding activity of CpTSP4 to HCT-8 cells. Therefore, CpTSP4 would contain additional motif(s) for binding to other ligand(s) on host cells that remains to be determined.

## Discussion

The early-branching *Cryptosporidium* is highly divergent from other apicomplexans in lifestyle (e.g., residing on top of host cells as an intracellular but extracytoplasmic parasite, rather than inside host cells) and metabolism (e.g., lacking an apicoplast, a typical mitochondrion and cytochrome-based respiratory chain) (37, 38). *Cryptosporidium* possesses two central microtubules with unknown biological roles, but lacks SPMT network (18). This study implies that the two *C. parvum* microtubules are involved in the anterograde trafficking of a micronemal protein and attached to γ-tubulin-based MTOC, which differ from SPMT in *Toxoplasma* and *Plasmodium* that use apical polar ring as “unconventional” MTOC to form a network to confer the elongated shape of the zoites (25, 39). The two central microtubules are unique to *Cryptosporidium*, as they are unseen in other apicomplexan lineages (e.g., coccidia and hematozoa). Therefore, their involvement in intracellular trafficking of a micronemal protein is not a common feature shared by apicomplexans.

Besides CpTSP4, only a limited number of other micronemal proteins have been experimentally confirmed by IFA and/or IEM for anterior location in sporozoites. These include CpGP900 (cgd7_4020), TRAPC1 (cgd1_3500), CpTSP8 (cgd6_780), CpROM1 (cgd3_980), cgd1_3550 (APPLE domain-containing protein), cgd2_1590 (APPLE/EGF-like domain-containing protein), and cgd7_1960 (WD40/YVTN repeat-like domain-containing protein) (9–11, 23). Among them, cgd7_1960 was more recently discovered as one of the proteins in the “microneme cluster” by HyperLOPIT technology and confirmed by IFA of hemagglutinin (HA)-tagged transgenic *C. parvum* (11). This HyperLOPIT-based study identifies several new candidate micronemal proteins based on their clustering with known microneme markers (e.g., CpGP900). It also clarifies that a previously suggested micronemal protein Cpa135 (cgd7_1730) is actually localized to the crystalloid body at the sporozoite posterior end. Nonetheless, these micronemal proteins show no filamentous distributions in sporozoites, suggesting that not all micronemal proteins utilize the two central microtubules for trafficking. This seems reasonable as many micronemal proteins are apically discharged.

The requirement of microtubules in the secretion of CpTSP4 is supported by IFA (distribution on the two microtubules) and by inhibition assay (complete inhibition of secretion by SB-743921 and SB-715992, the two highly selective kinesin-5/Eg5 inhibitors known for no off-target effect). Current data suggest anterograde transport of CpTSP4 in sporozoites, implying that CpTSP4 may be secreted from the posterior end and/or peripherally. However, it cannot be excluded that CpTSP4 may also be apically secreted, involving other types of microtubules that are unrecognizable by our anti-CpTubB antibody. Indeed, several short microtubule-like structures are observable on the apex of *C. parvum* sporozoites (40). The biological roles and potential involvement of these apical microtubules in the secretion of CpTSP4 remain to be elucidated. For comparison, *Toxoplasma* and *Plasmodium* contain short intraconoidal microtubules in their zoites that are implied for involvement in transport of vesicles (6, 8).

This study has also revealed a novel HBM responsible at least partially for the adhesive property of CpTSP4 to host cells. The other adhesive domain(s) in CpTSP4 and corresponding host ligand(s) remain to be determined. CpTSP4 lacks TMDs and is absent on the zoite plasma membranes as shown by IFA. Physiochemically, CpTSP4 is neutrally charged (pI = 7.45), thus unable to adhere to sporozoites or host cells by simple electrostatic interaction. Therefore, the released CpTSP4 would mostly adhere to the heparin/HS-containing mucus/ECM, but making no or little contribution to the adherence of the zoites. We speculate that secreted CpTSP4 adhering to the host cell mucus/ECM may serve as a molecular cushion to minimize direct contact of moving zoites to the intestinal mucins. This is beneficial to the moving zoites to avoid being restrained by the negatively charged mucins.

As globally distributed enteric parasites of medical and veterinary importance, only a single drug (nitazoxanide) is approved by FDA for treating cryptosporidiosis in immunocompetent patients. There is a lack of effective drug for use in immunocompromised patients (3, 41). Therefore, effective new anti-cryptosporidial drugs are needed. Kinesin-5/Eg5 has been an effective target for developing anticancer therapeutics and explored as anti-malarial target (42). This study shows that kinesin-5/Eg5 inhibitors could fully block the secretion of CpTSP4. Whether the two inhibitors, and their derivatives, could be explored/repurposed for developing anti-cryptosporidial drugs requires further investigations. However, CpKin5 shows sufficient sequence difference to human kinesin-5/Eg5 (73.3%/55.2% similarity/identity), suggesting that highly selective anti-CpKin5 inhibitors could be potentially developed for anti-cryptosporidial therapeutics or for use in studying the biology of kinesin-5 ortholog in the parasite.

## Materials and Methods

### Parasites and cell lines

A *C. parvum* isolate with gp60 subtype IIaA17G2R1 propagated in calves was used in all experiments. Oocysts were purified by sucrose gradient centrifugation protocols (43, 44), and stored in PBS containing 200 units/mL penicillin and 0.2 mg/mL streptomycin at 4°C. A human ileocecal epithelial cell line (HCT-8; ATCC # CCL-244) and bovine kidney cell (MDBK; ATCC # CCL-22) acquired from the Chinese Academy of Sciences Cell Bank (Shanghai) were cultured in RPMI-1640 medium containing 10% fetal bovine serum (FBS), 2 mM L-glutamine, 50 units/mL penicillin and 50 μg/mL streptomycin at 37°C in 5% CO_2_. For binding or infection assay, HCT-8 or MDBK cells were seeded into 24- or 48- or 96-well plates and allowed to grow overnight (∼80% confluence). For immunofluorescence assay (IFA), host cells were seeded into 48-well plates containing poly-L-lysine-treated round glass coverslips and infected with *C. parvum* by inoculation with oocysts or excysted sporozoites as specified. The invasion stage was prepared by infecting HCT-8 cell monolayers (∼60% confluence) with freshly excysted sporozoites for 15 to 30 min (9, 18, 23). Intracellular stages were prepared by inoculating HCT-8 monolayers with bleached and washed oocysts (in vitro excystation rate >80%) at 37°C for 3 h, followed by the removal of uninvaded parasites by medium exchange. Invaded parasites were allowed to grow for 3 to 72 h post-infection (hpi) or as specified.

### Antibody preparation

An anti-CpTSP4 mouse monoclonal antibody (mAb) was produced against a unique peptide (^89^KIKKADSWQEC^99^). Two specific pathogen-free mice were immunized five times with keyhole limpet hemocyanin (KLH)-linked peptide, including 100 μg with Freund’s complete adjuvant for first intradermal injection, 50 μg with incomplete adjuvant for next three intradermal injections, and 20 μg without adjuvant in final intravenous injection. Standard protocols were used in hybridoma production (45, 46). After three round of ELISA screening, the clone 9E6 was used for producing mouse ascites (anti-CpTSP4 mAb/clone 9E6).

The subtype of the mAb was IgG1 as determined using a subtype identification kit (Sigma-Aldrich, St. Louis, MO, USA). Non-immunoglobins in the ascites were removed by precipitation with caprylic acid, and immunoglobins were concentrated by ammonium sulfate precipitation (47). Purified mAb was stored at 2 μg/μL. Rabbit anti-CpGP900-C pAb and anti-CpTubB pAb were prepared and affinity-purified as described (23).

### Western blot analysis

*C. parvum* sporozoites were lysed in radio immunoprecipitation assay (RIPA) buffer containing protease inhibitor cocktail for mammals (Sigma-Aldrich). Sporozoite lysates (∼7.0×10^7^ per lane) in reducing sample buffer were heated at 95°C for 5 min, followed by fractionation in SDS-PAGE and transferred to nitrocellulose membranes. The blots were blocked in 5% BSA in TBST buffer (50 mM Tris-HCl at pH 7.5, 150 mM NaCl and 0.05% Tween-20) for 1 h and probed in 5% BSA-TBST buffer containing anti-CpTSP4 mAb (1:50 dilution) for 1 h. After three washes with TBST, blots were incubated with horseradish peroxidase (HRP)-conjugated goat anti-mouse IgG (Invitrogen, Waltham, MA, USA) and visualized using an enhanced chemiluminescence reagent (Beyotime Biotechnology, Shanghai, China). All procedures were conducted at room temperature unless specified.

### Indirect Immunofluorescence Assay (IFA)

Intact oocysts suspended in 4% paraformaldehyde were ruptured by three freeze-and-thaw cycles in liquid nitrogen and ice, followed by incubation for 30 min on ice and three washes with PBS. Free sporozoites were prepared by incubating oocysts in excystation medium (RPMI-1640 medium containing 0.75% taurodeoxycholic acid). Free merozoites were isolated from the supernatants of HCT-8 monolayers infected with *C. parvum* for 18–20 h. Intracellular parasites cultured on coverslips for various times were prepared as described above. Sporozoites and merozoites in suspension were fixed with cold 4% paraformaldehyde for 30 min and washed with PBS. Ruptured oocysts, sporozoites and merozoites after fixation were applied onto poly-L-lysine-coated glass slides. Samples were air-dried for 1 h, permeabilized with 0.2% Triton X-100 for 5 min and blocked for 1 h in 5% milk-PBS prepared from nonfat dry milk. Intracellular parasites on coverslips were similarly fixed, permeabilized and blocked. In gliding trail experiments, sporozoites after excystation were resuspended in RPMI-1640 medium, placed on poly-L-lysine-coated slides and incubated for 20 min at 37°C, followed by the same fixation, permeabilization and blocking procedures.

After blocking, specimens were incubated with specified antibodies in PBS containing 500 mM NaCl (high-salt PBS to minimize non-specific binding) for 1 h, washed three times with PBS, and incubated for 1 h with appropriate secondary antibodies conjugated with specified fluorophores (Thermo Scientific, West Palm Beach, FL), including Alexa Fluor 488-labeled donkey anti-mouse-IgG antibody, Alexa Fluor 594-labeled goat anti-rabbit-IgG or anti-mouse-IgG antibody. For invasion stage specimens, host cell F-actin was co-stained using rhodamine-phalloidin (Solarbio, Beijing, China). Samples were counter-stained for nuclei with 4,6-diamidino-2-phenylindole (DAPI, Sigma-Aldrich; 1.0 μg/mL). Procedures were performed at room temperature. Slides were examined under Olympus BX53 research fluorescence microscope. Images were capture with Olympus DP72 camera and stored in TIFF format. The relative fluorescence intensities were measured in some sporozoite samples and analyzed using ImageJ(Fiji). Images were processed with Adobe Photoshop (v2021 or higher), in which the signal levels might be linearly adjusted without local manipulations.

### ELISA detection of CpTSP4 secreted from sporozoites

The relative levels of CpTSP4 secreted from, and retained in, sporozoites during and after the excystation were detected by ELISA. Excystation was performed by incubating *C. parvum* oocysts (3×10^6^) in 75 μL excystation medium at 37°C for 45 min or as specified. In some assays, inhibitors (100 μM or as specified) were added into the excystation medium to evaluate their effects on the secretion of CpTSP4. Samples were centrifuged at 12,000 g for 5 min to collect supernatants and pellets. Some sporozoites after excystation and the removal of medium by centrifugation were further incubated in 75 μL RPMI-1640 medium at 37°C for 60 min, followed by centrifugation to collect supernatants and pellets. Sporozoite pellets were resuspended in 75 μL RPMI-1640 medium and subjected to five freeze-and-thaw cycles between liquid nitrogen and ice, followed by centrifugation for collecting supernatants. For comparison, intact oocysts (3×10^6^) suspended in 75 μL RPMI-1640 medium were subjected to the same freeze-and-thaw cycles and centrifugation to collect supernatants as described above. These supernatants were mixed with an equal volume of coating buffer (0.05 M carbonate-bicarbonate, pH 9.6) and all samples were calibrated to contain 0.375% sodium taurocholate.

Samples in coating buffer were added into 96-well plates in triplicates (50 μL/well, each equivalent to 10^6^ oocysts) and incubation overnight at 4°C. Plates were rinsed with 0.05% Tween-20/PBS and blocked with 5% milk/PBS (100 μL/well) at 37°C for 1 h. After three washes with 0.05% Tween-20/PBS, plates were incubated with anti-CpTSP4 mAb (2 μg/μL; 1:40 dilution) in high-salt PBS containing 500 mM NaCl (50 μL/well) 37°C for 1 h. For reference, a rabbit antiserum (1:40 dilution) against *C. parvum* total sporozoite proteins was used in parallel for all samples. After three washes with high-salt wash buffer, plates were incubated with alkaline phosphatase-conjugated goat anti-mouse IgG or anti-rabbit IgG(H+L) secondary antibodies (ImmunoWay Biotechnology, Plano, Texas; 1:10,000 dilution), followed by color development with p-nitrophenyl phosphate (Sigma-Aldrich). Optical density at 405 nm (OD_405_) was measured in a multifunctional microplate reader (BioTek, Vermont, USA).

### Heterologous expression of recombinant CpTSP4 and site-directed mutagenesis

CpTSP4 without signal peptide (cgd8_150; amino acids 27 to 484) was expressed as glutathione-S-transferase (GST)-fusion protein. Gene fragment was amplified from *C. parvum* genomic DNA by high-fidelity PCR using the following primers: CpTSP4-BamHӀ-F (5′-CGC<u>GGATCC</u>AAATACATTACCCCAGAAAAA-3′) and CpTSP4-EcoRӀ-R (5′-CCG<u>GAATTC</u>TTAGTTGAACAAATTTGGATCTA-3′) (underlines indicate restriction enzyme sites). Amplicons were cloned into a pGEX-4T-1 expression vectors (Invitrogen) at specified restriction sites. GST-tag alone was expressed using blank pGEX-4T-1 vector and used as negative control. In heparin-binding motif (HBM) validation assay, two Lys residues were replaced by Gln (K115Q/K116Q) by site-directed mutagenesis using the following primers in overlapping PCR: 1) CpTSP4-mut-BamHӀ-F1 (5′-GT<u>GGATCC</u>AAATACATTACCC**<u>C</u>**AG**<u>A</u>**AAAA-3′) and CpTSP4-mut-R1 (5′-ATAGCTCCATT**<u>G</u>**CT**<u>G</u>**ACACTTAATAAAGAAGAATGCA-3′); 2) CpTSP4-mut-F2 (5′-CTTTATTAAGTGT**<u>C</u>**AG**<u>C</u>**AATGGAGCTATGATGTAGAA-3′) and CpTSP4-EcoRӀ-R2 (5′-CCGG<u>GAATTC</u>TTAGTTGAACAAATTTGGATCTA-3′) (bold and underlines indicate mutation sites; underlines indicate restriction sites). All recombinant proteins were expressed in *Escherichia coli* strain BL21(DE3) cells and purified using glutathione-Sepharose 4B column following standard protocols. The purity and molecular weight were evaluated with SDS-PAGE.

### Host cell-binding assays

The host cell-binding of rCpTSP4 proteins (wild-type or mutated) was evaluated by immunofluorescence-based, flowcytometric and enzyme-linked assays. In immunofluorescence-based binding assay, HCT-8 cells were cultured to full confluence on coverslips placed in 48-well plates, fixed with 4% paraformaldehyde in PBS for 30 min on ice, blocked with 5% milk-PBS for 1 h, incubated with wild-type/mutated rCpTSP4 or GST-tag (each at 1.0 μM) in high-salt binding buffer (PBS containing 1.0 mM CaCl_2_, 1.0 mM MnCl_2_ and 500 mM NaCl) overnight at 4°C, fixed again with cold methanol for 10 min, and blocked again with 5% milk-PBS for 1 h. There were three washes in PBS after each step. Proteins bound to the host cells were detected using anti-GST mAb (ImmunoWay Biotechnology) at 37°C for 1 h, followed by incubation with Alexa Fluor 488-labeled Donkey anti-mouse-IgG antibody. Samples were examined under a fluorescence microscope as for IFA described above.

Flowcytometric assay was performed as described (48). Briefly, HCT-8 cells were cultured to semi-confluence at 37°C, washed twice with PBS, and treated with 500 μM EDTA in PBS for 15 min at 37°C. Detached cells (∼2×10^6^ in 400 μL) were washed with 2% FBS-PBS and incubated sequentially with 7.5 μM rCpTSP4 or GST in 2% FBS-PBS at 4°C for 2 h, anti-GST mAb (ABclonal Technology Co., Wuhan China) at 4°C for 1 h, and Alexa Fluor 488-labeled Donkey anti-mouse-IgG antibody at 1:1000 dilution for 4°C for 1 h. There were three washes with PBS after each step. At least 10,000 cells were analyzed with FACSCalibur system (BD Bioscience, Franklin Lakes, NJ, USA).

Enzyme-linked assay used previously described ELISA-like procedures (49–51). Briefly, HCT-8 cells were cultured to confluence in 96-well plates, fixed with 1% glutaraldehyde in PBS for 30 min, blocked with 5% milk-PBS for 1 h at 37°C, and treated with rCpTSP4 or GST-tag in 500 mM NaCl-PBS at specified concentrations overnight at 4°C. In some experiments, 20 μM heparin (Sigma-Aldrich, # H9399-5MU) or 20 μM chondroitin sulfate A (Sigma-Aldrich, # C9819-5G) was mixed with 2.0 μM rCpTSP4 (or GST-tag) in 500 mM NaCl/PBS, and then added to the fixed HCT-8 cells for 1 h at 37°C. Some HCT-8 cell specimens were treated with heparinase II (1.0 U/mL) (Sigma-Aldrich) in RPMI-1640 for 1 h at 37°C, followed by fixation with 1% glutaraldehyde for 30 min, three washes in PBS, and incubation with 1.0 μM rCpTSP4 or GST. Proteins on host cells was detected using ELISA procedures as described above.

### Microplate heparin-binding assays

Heparin-binding of CpTSP4 was evaluated by microplate-based and column/resin-based binding assays. Microplate assay resembles ELISA, in which 96-well plates were coated with heparin in 0.05 M carbonate-bicarbonate (pH 9.6) overnight at 4°C and blocked with 5% milk-PBS for 1 h at 37°C. In determining the binding kinetics, heparin-coated plates were incubated with serially diluted rCpTSP4 or rCpTSP4-mut (K115Q/K116Q), or GST-tag, in high-salt PBST buffer (500 mM NaCl and 1% Tween-20 in PBS) for 1 h at 37°C. In peptide competition assay, plates were incubated with 1.0 μM rCpTSP4 or GST-tag premixed with serially diluted synthetic peptides (HBM candidates) for 1 h at 37°C. The subsequent procedure followed those for standard ELISA as described above.

Column-based binding assay used heparin-agarose beads (GE Healthcare) as described (48, 52). Briefly, GST-CpTSP4 or GST-tag (1.0 μM or as specified) in PBS, with or without heparin or CSA (1.0 μM), was incubated with 40 μL heparin-agarose beads at 4°C for 2 h, washed five times with PBS, and boiled for 5 min in loading buffer. After quick centrifugation, supernatants were fractionated by SDS-PAGE, transferred onto nitrocellulose membrane, and probed with anti-GST antibody using standard western blot procedures.

### Evaluation of the binding kinetics of the p2-peptide to host cells and heparin

Heparin-coated microplates and formalin-fixed HCT-8 monolayers were prepared as described above. These microplates were blocked in 5% milk/PBS and incubated with FITC-conjugated p2-peptide (FITC-p2) at specified concentrations under the same experimental conditions as for assaying the binding of rCpTSP4. After washes with PBST buffer, relative fluorescence signals from FITC-p2 bound to heparin or host cells were detected using BioTek multifunctional microplate reader (*λ*_EX_ = 490 nm and *λ*_EM_ = 520 nm).

### Molecular docking

For binding of CpKin5 to SB-743921, CpKin5 model was retrieved from the AlphaFold Protein Structure Database (A3FPW8; https://alphafold.ebi.ac.uk/). The 3D-structure of the conserved N-terminal motor domain (amino acids from 1 to 371) showed high confidence (per-residue confidence score pLDDT > 90) was aligned, superimposed and analyzed using UCSF Chimera with a model of human Eg5–SB-743921 complex (PDB: 4BXN) (28). For heparin-CpTSP4 binding, a 51-aa region at the p2 site (^50^ASLCMDYVAF<u>FFIKC**KK**WSYDVEKKK</u>CLIKSGDRLCKYPD ENYISGLKNA S^101^; underline indicating HBM and bold fonts indicating critical residues) was extracted from the model (Q5CQ00). The heparin-binding was predicted using ClusPro 2.0 server under parameters specified for heparin-binding (33). Models were displayed using UCSF Chimera program (53).

### Statistical analysis

Quantitative experiments were performed three or more times independently with at least three biological replicates and two technical replicates. Charts and statistical analysis used GraphPad Prism (v9.0 or higher; San Diego, CA, USA). Statistical significances were evaluated by two-tailed Student’s *t*-test or two-way ANOVA with specified multiple comparisons.

### Animal studies

Animal experiments were approved by the Animal Welfare and Research Ethics Committee of Jilin University (AUP # 2020-1Z-20). Specific pathogen-free (SPF) female BALB/c mice (6 to 8-weeks old) and female rabbits were used in producing monoclonal or polyclonal antibodies. All animals were housed in an institutional approved facility.

## Supplemental Material

**Table S1.** List of TSP/TRAP family proteins in *Cryptosporidium parvum*.

**Table S2.** Similarity and identity scores of the motor domains between seven *C. parvum* kinesin. **Table S3.** Similarity and identity scores between kinesin-5 motor domains from selected species. **Fig. S1.** Additional images on immunostaining of CpTSP4 in *C. parvum* oocysts.

**Fig. S2.** Additional images of dual-labeling IFA of CpTSP4 with CpGP900 in sporozoites.

**Fig. S3.** Validation of antibody specificity in IFA for potential cross-reactions between CpTSP4 and CpTubB.

**Fig. S4.** Additional images on immunostaining of CpTSP4 (green) in gliding sporozoites.

**Fig. S5.** Additional images on immunostaining of CpTSP4 (green) in intracellular parasite.

**Fig. S6.** Additional images on immunostaining of CpTSP4 (green) in free merozoites.

**Fig. S7.** Additional images on the effect of the selective kinesin-5 inhibitor (SB723921) on the distribution of CpTSP4.

**Fig. S8.** Multiple sequence alignment of CpKin5 with orthologs from selected species.

**Fig. S9.** Molecular docking showing the binding of SB743921 to Cpkin5 motor domain that was superimposed with a human kinesin-5/Eg5 model (4BXN).

## Acknowledgements

This research was supported by grants of the National Key Research and Development Program of China (grant number 2022YFD1800200 to GZ), the National Natural Science Foundation of China (grant number 31772731 to JY) and National Key R&D Program of China (grant number 2017YFC1601200 to JY) and Jilin University Doctoral Interdisciplinary Research Fund (award number 101832020DJX094 to DQ). The funders had no role in the study design, data collection and data analysis, decision to publish, or preparation of the manuscript.

## References

1. Pumipuntu N, Piratae S. 2018. Cryptosporidiosis: A zoonotic disease concern. Vet World 11:681–686. doi:10.14202/vetworld.2018.681-686.

2. Khalil IA, Troeger C, Rao PC, Blacker BF, Brown A, Brewer TG, Colombara DV, De Hostos EL, Engmann C, Guerrant RL, Haque R, Houpt ER, Kang G, Korpe PS, Kotloff KL, Lima AAM, Petri WA, Jr., Platts-Mills JA, Shoultz DA, Forouzanfar MH, Hay SI, Reiner RC, Jr., Mokdad AH. 2018. Morbidity, mortality, and long-term consequences associated with diarrhoea from *Cryptosporidium* infection in children younger than 5 years: a meta-analyses study. Lancet Glob Health 6:e758–e768. doi:10.1016/S2214-109X(18)30283-3.

3. Checkley W, White AC, Jr., Jaganath D, Arrowood MJ, Chalmers RM, Chen XM, Fayer R, Griffiths JK, Guerrant RL, Hedstrom L, Huston CD, Kotloff KL, Kang G, Mead JR, Miller M, Petri WA, Jr., Priest JW, Roos DS, Striepen B, Thompson RC, Ward HD, Van Voorhis WA, Xiao L, Zhu G, Houpt ER. 2015. A review of the global burden, novel diagnostics, therapeutics, and vaccine targets for cryptosporidium. Lancet Infect Dis 15:85–94. doi:10.1016/S1473-3099(14)70772-8.

4. Dos Santos Pacheco N, Tosetti N, Koreny L, Waller RF, Soldati-Favre D. 2020. Evolution, Composition, Assembly, and Function of the Conoid in Apicomplexa. Trends Parasitol 36:688-704. doi:10.1016/j.pt.2020.05.001.

5. Tetley L, Brown SMA, McDonald V, Coombs GH. 1998. Ultrastructural analysis of the sporozoite of *Cryptosporidium parvum*. Microbiology (Reading) 144 (Pt 12):3249–3255. doi:10.1099/00221287-144-12-3249.

6. Cova MM, Lamarque MH, Lebrun M. 2022. How Apicomplexa Parasites Secrete and Build Their Invasion Machinery. Annu Rev Microbiol 76:619–640. doi:10.1146/annurev-micro-041320-021425.

7. Sharma P, Chitnis CE. 2013. Key molecular events during host cell invasion by Apicomplexan pathogens. Curr Opin Microbiol 16:432–7. doi:10.1016/j.mib.2013.07.004.

8. Dubois DJ, Soldati-Favre D. 2019. Biogenesis and secretion of micronemes in *Toxoplasma gondii*. Cell Microbiol 21:e13018. doi:10.1111/cmi.13018.

9. Gao X, Yin J, Wang D, Li X, Zhang Y, Wang C, Zhang Y, Zhu G. 2021. Discovery of New Microneme Proteins in *Cryptosporidium parvum* and Implication of the Roles of a Rhomboid Membrane Protein (CpROM1) in Host-Parasite Interaction. Front Vet Sci 8:778560. doi:10.3389/fvets.2021.778560.

10. Lendner M, Daugschies A. 2014. *Cryptosporidium* infections: molecular advances. Parasitology 141:1511–32. doi:10.1017/S0031182014000237.

11. Guerin A, Strelau KM, Barylyuk K, Wallbank BA, Berry L, Crook OM, Lilley KS, Waller RF, Striepen B. 2023. *Cryptosporidium* uses multiple distinct secretory organelles to interact with and modify its host cell. Cell Host Microbe 31:650–664 e6. doi:10.1016/j.chom.2023.03.001.

12. Kats LM, Cooke BM, Coppel RL, Black CG. 2008. Protein trafficking to apical organelles of malaria parasites - building an invasion machine. Traffic 9:176–86. doi:10.1111/j.1600-0854.2007.00681.x.

13. Sheiner L, Soldati-Favre D. 2008. Protein trafficking inside *Toxoplasma gondii*. Traffic 9:636–46. doi:10.1111/j.1600-0854.2008.00713.x.

14. Ravichandran Y, Goud B, Manneville JB. 2020. The Golgi apparatus and cell polarity: Roles of the cytoskeleton, the Golgi matrix, and Golgi membranes. Curr Opin Cell Biol 62:104–113. doi:10.1016/j.ceb.2019.10.003.

15. Barlan K, Gelfand VI. 2017. Microtubule-Based Transport and the Distribution, Tethering, and Organization of Organelles. Cold Spring Harb Perspect Biol 9. doi:10.1101/cshperspect.a025817.

16. Harding CR, Frischknecht F. 2020. The Riveting Cellular Structures of Apicomplexan Parasites. Trends Parasitol 36:979–991. doi:10.1016/j.pt.2020.09.001.

17. Venugopal K, Marion S. 2018. Secretory organelle trafficking in *Toxoplasma gondii*: A long story for a short travel. Int J Med Microbiol 308:751–760. doi:10.1016/j.ijmm.2018.07.007.

18. Wang C, Wang D, Nie J, Gao X, Yin J, Zhu G. 2021. Unique Tubulin-Based Structures in the Zoonotic Apicomplexan Parasite *Cryptosporidium parvum*. Microorganisms 9:1921. doi:10.3390/microorganisms9091921.

19. Deng M, Templeton TJ, London NR, Bauer C, Schroeder AA, Abrahamsen MS. 2002. *Cryptosporidium parvum* genes containing thrombospondin type 1 domains. Infect Immun 70:6987–95. doi:10.1128/IAI.70.12.6987-6995.2002.

20. John A, S MB, Madiedo Soler N, Wiradiputri K, Tichkule S, Smyth ST, Ralph SA, Jex AR, Scott NE, Tonkin CJ, Goddard-Borger ED. 2023. Conservation, abundance, glycosylation profile, and localization of the TSP protein family in *Cryptosporidium parvum*. J Biol Chem 299:103006. doi:10.1016/j.jbc.2023.103006.

21. Spano F, Putignani L, Naitza S, Puri C, Wright S, Crisanti A. 1998. Molecular cloning and expression analysis of a *Cryptosporidium parvum* gene encoding a new member of the thrombospondin family. Mol Biochem Parasitol 92:147–62

22. Putignani L, Possenti A, Cherchi S, Pozio E, Crisanti A, Spano F. 2008. The thrombospondin-related protein CpMIC1 (CpTSP8) belongs to the repertoire of micronemal proteins of *Cryptosporidium parvum*. Mol Biochem Parasitol 157:98–101. doi:10.1016/j.molbiopara.2007.09.004.

23. Li X, Yin J, Wang D, Gao X, Zhang Y, Wu M, Zhu G. 2022. The mucin-like, secretory type-I transmembrane glycoprotein GP900 in the apicomplexan *Cryptosporidium parvum* is cleaved in the secretory pathway and likely plays a lubrication role. Parasit Vectors 15:170. doi:10.1186/s13071-022-05286-8.

24. Barnes DA, Bonnin A, Huang JX, Gousset L, Wu J, Gut J, Doyle P, Dubremetz JF, Ward H, Petersen C. 1998. A novel multi-domain mucin-like glycoprotein of *Cryptosporidium parvum* mediates invasion. Mol Biochem Parasitol 96:93–110. doi:10.1016/s0166-6851(98)00119-4.

25. Tosetti N, Dos Santos Pacheco N, Bertiaux E, Maco B, Bournonville L, Hamel V, Guichard P, Soldati-Favre D. 2020. Essential function of the alveolin network in the subpellicular microtubules and conoid assembly in *Toxoplasma gondii*. Elife 9. doi:10.7554/eLife.56635.

26. Mogre SS, Brown AI, Koslover EF. 2020. Getting around the cell: physical transport in the intracellular world. Phys Biol 17:061003. doi:10.1088/1478-3975/aba5e5.

27. Myers SM, Collins I. 2016. Recent findings and future directions for interpolar mitotic kinesin inhibitors in cancer therapy. Future Med Chem 8:463–89. doi:10.4155/fmc.16.5.

28. Talapatra SK, Anthony NG, Mackay SP, Kozielski F. 2013. Mitotic kinesin Eg5 overcomes inhibition to the phase I/II clinical candidate SB743921 by an allosteric resistance mechanism. J Med Chem 56:6317–29. doi:10.1021/jm4006274.

29. Zeeshan M, Shilliday F, Liu T, Abel S, Mourier T, Ferguson DJP, Rea E, Stanway RR, Roques M, Williams D, Daniel E, Brady D, Roberts AJ, Holder AA, Pain A, Le Roch KG, Moores CA, Tewari R. 2019. *Plasmodium* kinesin-8X associates with mitotic spindles and is essential for oocyst development during parasite proliferation and transmission. PLoS Pathog 15:e1008048. doi:10.1371/journal.ppat.1008048.

30. Rudd TR, Preston MD, Yates EA. 2017. The nature of the conserved basic amino acid sequences found among 437 heparin binding proteins determined by network analysis. Mol Biosyst 13:852–865. doi:10.1039/c6mb00857g.

31. Torrent M, Nogues MV, Andreu D, Boix E. 2012. The “CPC clip motif”: a conserved structural signature for heparin-binding proteins. PLoS One 7:e42692. doi:10.1371/journal.pone.0042692.

32. Jumper J, Evans R, Pritzel A, Green T, Figurnov M, Ronneberger O, Tunyasuvunakool K, Bates R, Zidek A, Potapenko A, Bridgland A, Meyer C, Kohl SAA, Ballard AJ, Cowie A, Romera-Paredes B, Nikolov S, Jain R, Adler J, Back T, Petersen S, Reiman D, Clancy E, Zielinski M, Steinegger M, Pacholska M, Berghammer T, Bodenstein S, Silver D, Vinyals O, Senior AW, Kavukcuoglu K, Kohli P, Hassabis D. 2021. Highly accurate protein structure prediction with AlphaFold. Nature 596:583–589. doi:10.1038/s41586-021-03819-2.

33. Mottarella SE, Beglov D, Beglova N, Nugent MA, Kozakov D, Vajda S. 2014. Docking server for the identification of heparin binding sites on proteins. J Chem Inf Model 54:2068–78. doi:10.1021/ci500115j.

34. Azzouz N, Kamena F, Laurino P, Kikkeri R, Mercier C, Cesbron-Delauw MF, Dubremetz JF, De Cola L, Seeberger PH. 2013. *Toxoplasma gondii* secretory proteins bind to sulfated heparin structures. Glycobiology 23:106–20. doi:10.1093/glycob/cws134.

35. Carruthers VB, Hakansson S, Giddings OK, Sibley LD. 2000. *Toxoplasma gondii* uses sulfated proteoglycans for substrate and host cell attachment. Infect Immun 68:4005–11. doi:10.1128/IAI.68.7.4005-4011.2000.

36. Bambino-Medeiros R, Oliveira FO, Calvet CM, Vicente D, Toma L, Krieger MA, Meirelles MN, Pereira MC. 2011. Involvement of host cell heparan sulfate proteoglycan in *Trypanosoma cruzi* amastigote attachment and invasion. Parasitology 138:593–601. doi:10.1017/S0031182010001678.

37. Templeton TJ, Enomoto S, Chen WJ, Huang CG, Lancto CA, Abrahamsen MS, Zhu G. 2010. A genome-sequence survey for Ascogregarina taiwanensis supports evolutionary affiliation but metabolic diversity between a *Gregarine* and *Cryptosporidium*. Mol Biol Evol 27:235–48. doi:10.1093/molbev/msp226.

38. Rider SD, Jr., Zhu G. 2010. *Cryptosporidium*: genomic and biochemical features. Exp Parasitol 124:2–9. doi:10.1016/j.exppara.2008.12.014.

39. Striepen B, Jordan CN, Reiff S, van Dooren GG. 2007. Building the perfect parasite: cell division in apicomplexa. PLoS Pathog 3:e78. doi:10.1371/journal.ppat.0030078.

40. Mageswaran SK, Guerin A, Theveny LM, Chen WD, Martinez M, Lebrun M, Striepen B, Chang YW. 2021. In situ ultrastructures of two evolutionarily distant apicomplexan rhoptry secretion systems. Nat Commun 12:4983. doi:10.1038/s41467-021-25309-9.

41. Schneider A, Wendt S, Lubbert C, Trawinski H. 2021. Current pharmacotherapy of cryptosporidiosis: an update of the state-of-the-art. Expert Opin Pharmacother doi:10.1080/14656566.2021.1957097:1-6. doi:10.1080/14656566.2021.1957097.

42. Liu L, Richard J, Kim S, Wojcik EJ. 2014. Small molecule screen for candidate antimalarials targeting Plasmodium Kinesin-5. J Biol Chem 289:16601–14. doi:10.1074/jbc.M114.551408.

43. Truong Q, Ferrari BC. 2006. Quantitative and qualitative comparisons of *Cryptosporidium* faecal purification procedures for the isolation of oocysts suitable for proteomic analysis. Int J Parasitol 36:811–9. doi:10.1016/j.ijpara.2006.02.023.

44. Zhang H, Zhu G. 2020. High-Throughput Screening of Drugs Against the Growth of *Cryptosporidium parvum* In Vitro by qRT-PCR. Methods Mol Biol 2052:319–334. doi:10.1007/978-1-4939-9748-0_18.

45. Hadavi R, Zarnani AH, Ahmadvand N, Mahmoudi AR, Bayat AA, Mahmoudian J, Sadeghi MR, Soltanghoraee H, Akhondi MM, Tarahomi M, Jeddi-Tehrani M, Rabbani H. 2010. Production of Monoclonal Antibody against Human Nestin. Avicenna J Med Biotechnol 2:69–77

46. Greenfield EA. 2018. Polyethylene Glycol Fusion for Hybridoma Production. Cold Spring Harb Protoc 2018. doi:10.1101/pdb.prot103176.

47. McKinney MM, Parkinson A. 1987. A simple, non-chromatographic procedure to purify immunoglobulins from serum and ascites fluid. J Immunol Methods 96:271–8. doi:10.1016/0022-1759(87)90324-3.

48. Inomata A, Murakoshi F, Ishiwa A, Takano R, Takemae H, Sugi T, Cagayat Recuenco F, Horimoto T, Kato K. 2015. Heparin interacts with elongation factor 1alpha of *Cryptosporidium parvum* and inhibits invasion. Sci Rep 5:11599. doi:10.1038/srep11599.

49. Chatterjee A, Banerjee S, Steffen M, O’Connor RM, Ward HD, Robbins PW, Samuelson J. 2010. Evidence for mucin-like glycoproteins that tether sporozoites of *Cryptosporidium parvum* to the inner surface of the oocyst wall. Eukaryot Cell 9:84–96. doi:10.1128/EC.00288-09.

50. Bhat N, Joe A, PereiraPerrin M, Ward HD. 2007. *Cryptosporidium* p30, a galactose/N-acetylgalactosamine-specific lectin, mediates infection in vitro. J Biol Chem 282:34877–87. doi:10.1074/jbc.M706950200.

51. Bhalchandra S, Ludington J, Coppens I, Ward HD. 2013. Identification and characterization of *Cryptosporidium parvum* Clec, a novel C-type lectin domain-containing mucin-like glycoprotein. Infect Immun 81:3356–65. doi:10.1128/IAI.00436-13.

52. Zhang Y, Jiang N, Lu H, Hou N, Piao X, Cai P, Yin J, Wahlgren M, Chen Q. 2013. Proteomic analysis of *Plasmodium falciparum* schizonts reveals heparin-binding merozoite proteins. J Proteome Res 12:2185–93. doi:10.1021/pr400038j.

53. Pettersen EF, Goddard TD, Huang CC, Meng EC, Couch GS, Croll TI, Morris JH, Ferrin TE. 2021. UCSF ChimeraX: Structure visualization for researchers, educators, and developers. Protein Sci 30:70–82. doi:10.1002/pro.3943.

